# *In vitro* and *in vivo* validation of recombinant Syrian hamster interferon-α

**DOI:** 10.64898/2026.11.12.766184

**Authors:** Victor Neven, Steven Beelen, Elias Broeckhoven, Brigitta M. Laksono, Sergei V. Strelkov, Yeranddy A. Alpizar, Kai Dallmeier

## Abstract

The Syrian hamster (*Mesocricetus auratus*) is an important model for the study of viral pathogenesis and the development of antiviral drugs and vaccines, yet species-specific reagents remain scarce. Here, we validate the potent and specific activity of recombinantly expressed Syrian hamster interferon-α-12-like protein (maIFNα) *in vitro* and *in vivo*. MaIFNα expressed in *E. coli* induced *MX2* and *ISG15* expression in both BHK21 hamster kidney cells as well as in primary hamster fibroblasts in a dose-dependent manner, with EC_50_ values in low pg/mL range. Activation was strictly STAT2-dependent and more potent, by at least one order of magnitude, than that achieved by heterologous human or mouse IFNα. ISG induction translated into a strong antiviral activity against YF17D virus replication in BHK21 cells as well as ISRE-dependent reporter expression in a newly established hamster reporter cell line. When tested in wild-type but not in STAT2-deficient hamsters, maIFNα showed a physiologic, vigorous yet rapidly contracting systemic ISG response. Together these findings establish maIFNα as a functional antiviral cytokine and valuable reagent for preclinical infectious disease research using Syrian hamsters.

## INTRODUCTION

Type I interferons (IFNs) are cytokines encoded by a group of intron-less, single exon genes conserved among vertebrates (1). They constitute a key early component of the antiviral response by inducing a cellular antiviral state that restricts viral replication and spread, while activating innate and adaptive immunity (2). In humans, type I IFNs comprise 16 subtypes across five classes (i.e. 12 IFNα subtypes, IFNβ, IFNω, IFNε and IFNκ), which share a common structural fold consisting of four parallel α-helices despite a considerable sequence diversity of as little as 35% to 95% amino acid (AA) identity (3). The IFNα class accounts for 12 subtypes with at least 76% sequence identity encoded by 13 *IFNA* genes (4). Cellular expression of IFNs is conditional and subtype-specific: IFNβ is expressed by virtually all nucleated cells whereas IFNα is mainly expressed by plasmacytoid dendritic cells (5). IFNω is primarily expressed by leukocytes (6), and expression of IFNε and IFNκ is restricted to mucosal tissues (7) and keratinocytes (8), respectively. Expression of IFNs is triggered by environmental stimuli such as viral infections sensed by pathogen recognition receptor (PPR)-mediated pathways (9).

All type I IFNs bind the same ubiquitously expressed heterodimeric transmembrane IFNα receptor (IFNAR), composed of the IFNAR1 and IFNAR2 subunits. Intracellular signaling proceeds via the receptor-associated protein tyrosine kinases Janus kinase 1 (JAK1) and tyrosine kinase 2 (TYK2), which phosphorylates the cytoplasmic transcription factors Signal Transducer and Activator of Transcription 1 (STAT1) and STAT2, the resulting heterodimer translocates to the nucleus and assembles with IFN-regulatory factor 9 (IRF9) into the IFN-stimulated gene factor 3 (ISGF3) complex, which binds to the IFN-stimulated response element (ISRE) upstream of a large number of IFN-stimulated genes (ISG). ISG expression results in the inhibition of viral transcription, translation and replication, the degradation of viral nucleic acids and the alteration of cellular lipid metabolism (2,10). To cope with this broad innate antiviral response, several viruses encode numerous antagonists of the IFN pathway (11,12) which permits their replication and, in the absence of an effective adaptive immune response, progression to severe viral disease.

Antiviral drugs and vaccines are essential for protection against existing and emerging viral diseases, and their development continues to rely on predictive animal models. Among these, the Syrian hamster (*Mesocricetus auratus*) has emerged as a valuable preclinical model because it faithfully reproduces key pathological and clinical features of human infections caused by a diverse range of RNA and DNA viruses (orthoflavi-, alpha-, corona-, orthobunya-, phlebo-, arena-, paramyxo-, filo- and adenoviruses) (13–16) , including several important (re-)emerging pathogens (e.g. yellow fever virus (17) and Oropouche virus (18)) and WHO Priority Pathogens (e.g., SARS-CoV-2 (19), Ebola virus (20) and Nipah virus (21)). However, the availability of thoroughly functionally validated, species-specific immunological reagents for hamsters remains limited.

Hamster IFNα (maIFNα) has only been partially characterized (22–24). A current genome assembly (BCM_Maur_2.0, GCF_017639785.1) contains a cluster of 14 maIFNα genes predicted to encode functional proteins plus five predicted pseudogenes; twelve of the former annotated as maIFNα-12-like (190 AA) and two as maIFNα-9-like (182 AA) (25). Experimental confirmation is lacking and currently no bioactive maIFNα is available, constraining assay development and essentially limiting experimental assessment of IFNα responses in this species. Here, we recombinantly express the predicted maIFNα-12-like protein (hereafter maIFNα) and characterize its biological function regarding its antiviral activity and ISG induction *in vitro* and *in vivo*.

## MATERIALS AND METHODS

### Sequence alignment and protein structure prediction

Annotated and predicted protein sequences were retrieved from NCBI protein (https://www.ncbi.nlm.nih.gov/protein/) (**Table 1**) and aligned omitting the respective signal peptide sequences using MEGA12 software (https://www.megasoftware.net/, version 12.1.2 (26)). ClustalW was used with default parameters and Gonnet protein weight matrix. Phylogenetic tree was constructed by Neighbor-Joining method (27) using MEGA12 software (Jones-Taylor-Thornton (28)). Sequence conservation statistics were obtained with Clustal Omega tool from EMBL-EBI (https://www.ebi.ac.uk/jdispatcher/msa/clustalo) with default parameters. Three-dimensional structures of human IFNα-2a (hsIFNα-2a), mouse IFNα-12 (mmIFNα-12) and maIFNα were predicted using AlphaFold server 3 (Google DeepMind, https://alphafoldserver.com/ (29)) and annotated using Mol* 3D viewer (RCSB PDB, https://www.rcsb.org/3d-view). Predictions were obtained with predicted aligned errors (PAE). Confidences scores were given as plDDT values.

**Table 1.** Species origin, gene products and NCBI accession numbers of IFNα proteins.

| **Species** | **Gene product** | **NCBI accession number** |
|---|---|---|
| *Homo sapiens* (hs) | IFNα-2a (hsIFNα-2a) | NP_000596.2 |
| *Mus musculus* (mm) | IFNα-9 (mmIFNα-9*)* | NP_034637.1 |
| *Mus musculus* (mm) | IFNα-12 (mmIFNα-12*)* | AAI04353.1 |
| *Mesocricetus auratus* (ma) | IFNα-9-like (maIFNα-9-like) | XP_040613376.1 |
| *Mesocricetus auratus* (ma) | IFNα-12-like (maIFNα) | XP_040613374.1 |
| *Cricetulus griseus* (cg) | IFNα-9 (cgIFNα-9) | XP_027256146.1 |
| *Cricetulus griseus* (cg) | IFNα-12 (cgIFNα-12) | XP_027256148.1 |
| *Phodopus roborovskii* (pr) | IFNα-9-like (prIFNα-9-like) | XP_051043925.1 |
| *Phodopus roborovskii* (pr) | IFNα-12-like (prIFNα-12) | XP_051043952.1 |
| *Rattus norvegicus* (rn) | IFNα-12-like (rnIFNα-12) | XP_038967487 |
| *Gallus gallus* (gg) | IFNα (ggIFNα) | ABY73738.1 |
| *Danio rerio* (dr) | Interferon (drIFN) | AAM95448.1 |

### Cloning and bacterial protein expression

The gene sequence of the predicted maIFNα protein was codon optimized for expression in *E. coli* using the Integrated DNA Technologies (IDT) Codon Optimization Tool (https://eu.idtdna.com/pages/tools/codon-optimization-tool) and synthesized, without signal peptide and stop codon, by IDT. The gene fragment was cloned in the pETSUK2 plasmid (30), encoding an N-terminal hexahistidine (His_6_)-tag followed by the small ubiquitin-like modifier (SUMO), via homologous recombination using the NEBuilder® HiFi DNA Assembly Cloning Kit (New England Biolabs, Ipswich, USA), following linearization of the plasmid with HindIII (New England Biolabs) and KpnI (New England Biolabs) restriction enzymes.

The resulting pETSUK2-maIFNα plasmid was transformed in *E. coli* Rosetta 2 (DE3) pLysS strain (Merck, Rahway, USA) and grown overnight (ON) at 37 °C on LB agar plate containing 100 µg/mL ampicillin (Acros, Geel, Belgium) and 30 µg/mL chloramphenicol (Roth, Newport Beach, USA).

The cells were grown in ZYP-5052 auto-inducing medium (31,32) containing 100 µg/mL ampicillin and 30 µg/mL chloramphenicol at 24 °C until absorbance at 600 nm reached a value of 4. The culture was further incubated for 24 h at 18 °C after which the cells were collected following centrifugation (45 min, 12,000 x *g*, 4 °C, rotor #12269, Biosafe, Sigma, Kawasaki, Japan) and stored at -80 °C until further use.

### Protein purification

The protein was purified using a previously reported protocol (33). The bacterial pellet (∼7 g) was thawed on ice and resuspended in 100 mL low imidazole buffer (50 mM sodium phosphate pH 7.5 (Sigma), 250 mM NaCl (Acros), 12.5 mM imidazole pH 7.5 (Acros), 5 mM β-mercaptoethanol (Sigma) with Sigma Fast protease inhibitor (Sigma Aldrich, St. Louis, USA) and 10 mM MgCl_2_ (Acros)). The cells were lysed by sonication on ice at 50% amplitude with 2 s on / 4 s off pulses for a total sonication time of 3 min, repeated three times with 10 min incubation between cycles (Branson Digital Sonifier). The lysate was centrifuged (1 h, 16,000 x *g,* 4 °C, rotor #12500, Sigma 6K15) to collect supernatant.

Supernatant was applied on a HIS-Select® Nickel (Ni) Affinity Gel column (Merck), pre-equilibrated with low imidazole buffer. The column was washed with four column volumes (CV) low imidazole buffer and 0.1% Triton X-100 (Fisher Scientific, Pittsburgh, USA), followed by a second wash step with 2 CV low imidazole buffer. The target protein was eluted with high imidazole buffer (50 mM sodium phosphate pH 7.5, 250 mM NaCl, 500 mM imidazole pH 7.5) and His_6_-SUMO tag was removed with His_6_-tagged SUMO hydrolase (34) (ratio 1:1000) during ON dialysis at 4 °C to exchange high imidazole buffer with low imidazole buffer. The sample was re-applied on the Ni-chelating column to capture uncleaved protein, the cleaved tag and the SUMO hydrolyase. The untagged maIFNα was collected in the flow-through and dialyzed ON at 4 °C to the ion exchange buffer (20 mM Tris-HCl (Acros), 2 mM DTT (Acros), 10% w/v glycerol v, pH 7.5). The sample was then applied onto two consecutive 5 mL HiTrap Q HP anion-exchange columns (Cytiva, Chicago, USA, 10 mL total column volume) and equilibrated with 20 mM Tris-HCL pH 7.5, 2.5 mM DTT, using the Äkta™ Pure system with UV detector (Cytiva). The protein was eluted with a linear gradient of NaCl of 40% with a high salt buffer (20 mM Tris-HCL PH7.5, 2.5 mM DTT, 1 M NaCl). The eluted fractions (4 mL) containing the target protein were pooled after SDS-PAGE analysis, concentrated to 2 mL using a 10 kD cut-off filter (Merck) and applied onto a Superdex 75 16/600 PG column (Cytiva) equilibrated with size exclusion buffer (20 mM Tris-HCl pH 7.5, 2.5 mM DTT, 150 mM NaCl, pH 7.5). Fractions containing the pure target protein were pooled after SDS-PAGE analysis, aliquoted and flash-frozen before being stored at -80 °C until further use. Absence of endotoxin was confirmed by *Limulus* amebocyte lysate (LAL) assay (ToxinSensor™ Chromogenic LAL Endotoxin Assay Kit, GenScript, Piscataway, USA) yielding undetectable LPS levels (< 0.1 EU/µg protein).

### SDS-PAGE analysis

Collected fractions and all products from each purification step were analyzed using SDS-PAGE (17%). Collected fractions (10 µL) were added to 20 µL ddH2O and 10 µL loading buffer (0.25 M Tris-HCl pH 6.8, 40% glycerol, 0.28 M SDS, 0.2% Bromophenol Blue) with 0.14 M β-mercaptoethanol and boiled for 5 min at 95 °C. The sample (7 µL) is loaded on a 17% SDS-Page gel and ran using Laemmli buffer (25 mM Trizma Base, 200 mM glycine, 0.1 w/v% SDS, pH 8.6). The gel was submerged in fixation buffer (40% ethanol, 10% acetic acid) for 10 min and stained with Coomassie Blue (25% ethanol, 8% acetic acid, 0.12% Coomassie Blue) until it was homogenously stained (30 min). The gel was subsequently incubated in destaining buffer (25% ethanol, 8% acetic acid) for 30 min and allowed to fully de-stain ON.

### Cells

Baby hamster kidney fibroblast (BHK21J, kindly provided by P. Bredenbeek, Leiden University Medical Centre, the Netherlands) cells were maintained in Modified Eagle Medium (MEM; Gibco, Waltham, USA) containing 10% fetal calf serum (FCS; Gibco), 1% penicillin–streptomycin-glutamine (Pen Strep; Gibco) and 1% non-essential amino acids (NEAA; Gibco). HEK-Blue™ ISG cells (hkb-isg-1, Invivogen, San Diego, USA), B16-Blue™ IFN-α/β cells (bb-ifnt1, Invivogen) and BHK-miRFP670 cells (constitutively expressing near-infrared fluorescent protein miRFP670 as translational fusion to histone protein H2B and generated based on previously established protocols in-house (35)) were maintained in Dulbecco’s Modified Eagle Medium (DMEM, Gibco) containing 10% FCS (Gibco) and 1% Pen Strep (Gibco). HEK-Blue™ ISG and B16-Blue™ IFN-α/β cells were selected with 100 µg/mL Zeocin (Invivogen, ant-zn-1). BHK-miRFP670 cells were selected with 10 µg/mL blasticidin (Invivogen, ant-bl-1). All cells were incubated at 37 °C with 5% CO_2_.

### Stable reporter cell line generation

The mNeonGreen (mNG) DNA sequence was cloned into pLenti-EFIα-H2B-miRFP670-P2A-Blast plasmid (35) (containing the coding sequence for the bright monomeric near-infrared fluorescent protein (miRFP670 (36)) with a 5’ nuclear H2BC11 histone tag). A synthetic IFN-inducible ISRE promotor, containing a 5x ISRE repeat sequence, was cloned together with the mNG sequence upstream of the EFI-α promotor via homologous recombination using the NEBuilder® HiFi DNA Assembly Cloning Kit (New England Biolabs). The resulting pLenti-ISRE-mNG-H2B-miRFP670-P2A-Blast vector was used to produce lentiviral particles, following a previously established protocol (37). Briefly, HEK-AD cells were transfected using the TransIT®-LT1 Transfection Reagent (Mirus Bio, Madison, USA) with the pLenti-ISRE-mNG-H2B-miRFP670-P2A-Blast plasmid and lentiviral packaging plasmids psPAX2 (packaging plasmid; Addgene, #12260) and pMD2.G (VSV-G envelop plasmid; Addgene, #12259) in a 1:1:1 molar ratio. Supernatant containing lentiviral particles was collected at 2 days post-transfection (dpt). Next, BHK21J cells were stably transfected with the lentivirus. From 5 days onward, stably transduced cells BHK-ISRE-mNG cells were selected with 30 µg/mL blasticidin.

To validate the reporter assay, BHK-ISRE-mNG cells were seeded (65,000 cells per well) in a 48-well plate (Corning, New York, USA) in 180 µL MEM medium containing 2% FCS. A 10x serial dilution (20 µL in 2% FCS medium) of maIFNα or Alfacon (38) (Universal^TM^ Type I IFN Human IFN-Alpha Hybrid Protein, PBL Assay Science, Piscataway Township, USA) was added to the cells directly after seeding. After 24 h, cells were collected, transferred to a U-bottom plate (Corning) and washed with FACS buffer (PBS containing 2% FCS). After washing, the cells were fixed with 4% paraformaldehyde (PFA) for 10 min and resuspended in 200 µL FACS buffer. Expression of mNG and miRFP670 was determined using a FACS Canto^TM^ II (BD Biosciences, Franklin Lakes, USA). Data were processed in FlowJo (for gating strategy see **Supplementary Figure S2C**).

### Primary fibroblasts isolation

Primary hamster fibroblasts were isolated and maintained according to previously established protocol (39). Briefly, ears from wild-type (WT) and STAT2-knockout (KO) hamsters were shaved and diced using a scalpel. Ear fragments from 1 ear were cultured in a T75-flask with 5 mL initial growth medium (DMEM, 20% FCS, 1% Pen Strep (Gibco), 1% L-glutamine (Gibco), 1% amphotericin B (Gibco), 0.5% gentamicin sulfate (Thermo Fisher Scientific), 100 µg/mL normocin (Invivogen)) for 4 days. The medium was removed and replaced with 5 mL initial growth medium at days 4, 7, 9 and 11. After 14 days, the cells were passaged for the first time: the medium was removed, cells were washed twice with HBSS (Gibco) and disassociated with 2 mL trypsin 0.05% (Gibco). Cells were resuspended in 8 mL expansion growth medium (DMEM, 20% FCS, 1% Pen Strep, 1% L-glutamine, 1% amphotericin B, 0.5% gentamicin sulfate) and centrifuged (5 min, 500 x *g*, room temperature, rotor S-4-104, Eppendorf, Hamburg, Germany). The cells were resuspended in 9 mL expansion growth medium and filtered through a 70-µm cell strainer. Cells were transferred to a T75-flask after filtration. Subsequent passaging was performed as before, excluding the cell filtering. Cells were incubated at 37 °C with 5% CO_2_ (**Supplementary Figure S2A**).

### *In vitro* characterization of maIFNα

BHK21J cells and primary fibroblasts (WT and STAT2-KO) were seed (60,000 cells per well) 24 h before the assay in a 48-well plate (Corning) in MEM or DMEM respectively, containing 10% FCS. The day after, the medium was replaced with 2% FCS medium containing a serial dilution of purified maIFNα protein. As a positive control, cells were stimulated with Alfacon. Cells were lysed 8 h post-treatment and RNA was extracted following the RNeasy Mini protocol (Qiagen, Venlo, the Netherlands). RNA levels of *MX2* (GenBank: EU616539.1)*, ISG15* (NCBI RefSec: XM_013119951.3) and β*-ACTIN* (NCBI RefSec: NM_001281595.1) were determined using the iTaq Universal Probes One-Step Kit (BioRad, Hercules, USA) on the LightCycler® 96 system (Roche Diagnostics, Indianapolis, USA) following the manufacturer’s protocol. The reverse transcriptase step was carried out for 10 min at 55 °C. PCR was performed for 3 min at 95 °C, followed by 45 cycles of 15 s at 95 °C and 30 s at 55 °C. Primers and probes (**Table 2**) were synthesized by IDT. The median of housekeeping gene *β-ACTIN* was used for normalization. The relative fold change was calculated using the 2^−ΔΔCt^ method (40).

**Table 2.**
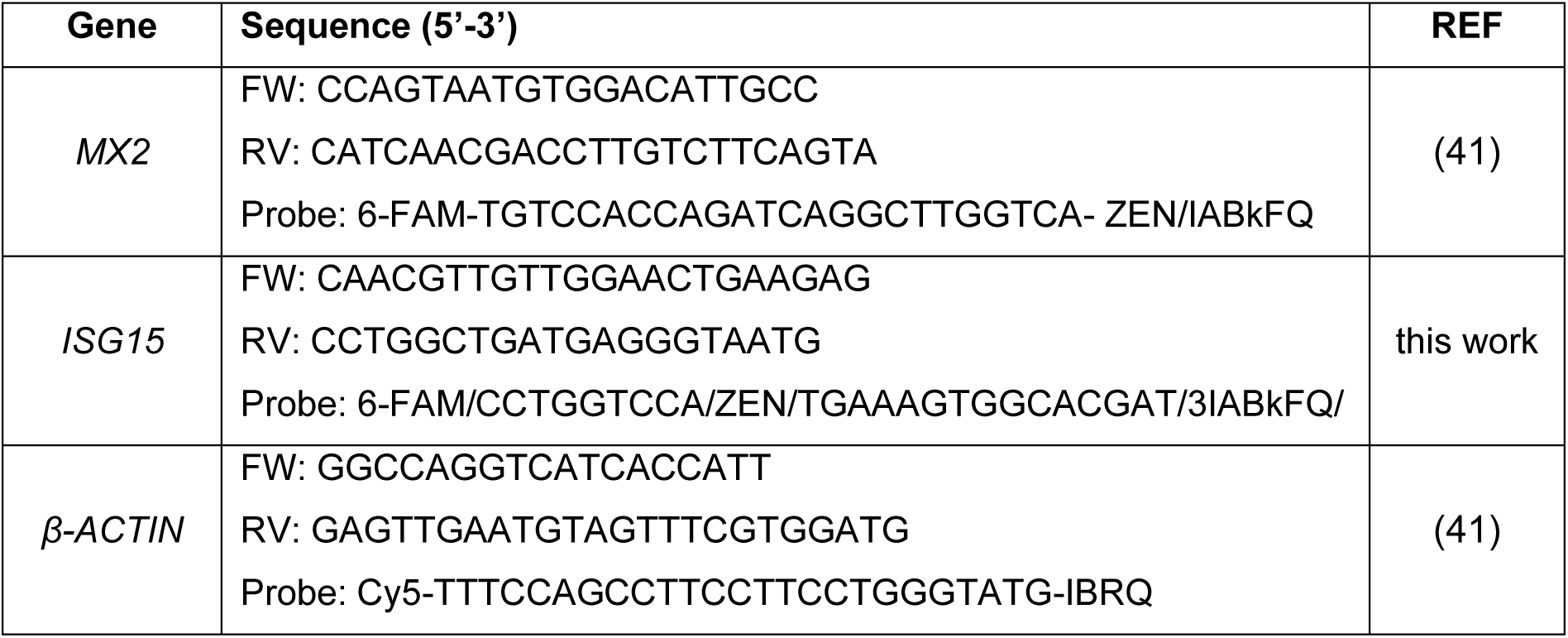
List of primers used for qPCR analyses.

### Antiviral assay

BHK-miRFP670 cells were seeded (15,000 cells per well) in a black 96-well plate (Greiner Bio-One, Kremsmünster, Austria) in DMEM containing 2% FCS and incubated ON at 37 °C with 5% CO_2_. Medium was removed the day after and 100 µL of a serial dilution of purified maIFNα, hsIFNα-2a, mmIFNα-1 or Alfacon in medium containing 2% FCS was added. After 1 h and without removal of the respective IFNs, cells were inoculated with a multiplicity of infection (MOI) of 0.01 of yellow fever vaccine-based YF17D-mCherry reporter virus (42) (100 µL, 75 PFU, in medium containing 2% FCS) and incubated at 37 °C with 5% CO_2_ for 9 days. High content image acquisition was performed using an Operetta CLS High Content Imaging and Analysis system (Revvity, Inc., Waltham, Massachusetts, USA). Each day, 9 images per well were taken in non-confocal mode with a 10x objective. For total cell counts, BHK-miRFP670 reporter (miRFP670^+^) cells were excited with a 615-645 LED source and emission was detected with a 655-760 nm filter. For detection of the YF17D-mCherry reporter virus-infected (miRFP670^+^ mCherry^+^) cells, cells were excited with a 530-560 LED source, and emission was detected with a 570-650 nm filter. % inhibition of viral replication was calculated as:

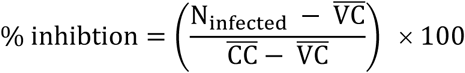

Where N_infected_ is the number of infected cells (miRFP670^+^ mCherry^+^), 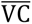 is the average of infected cells in the virus treated controls and 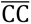 is the average of non-infected cells (miRFP670^+^ mCherry^−^) in the untreated cell controls.

Viability of the cells following IFN stimulation was calculated as:

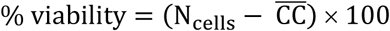

Where N_cells_ is the respective number of miRFP670^+^ cells and 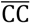 is the average of miRFP670^+^ cells in the cell controls.

### Cross-species interferon activity assay

HEK-Blue™ ISG cells and B16-Blue™ IFN-α/β cells, expressing a secreted embryonic alkaline phosphatase (SEAP) in response to IFN stimulation, were seeded in 96-well plates (Corning) at 35,000 and 75,000 cells per well, respectively, in 180 µL medium containing 2% FCS. A 10x serial dilution of hsIFNα-2a, mmIFNα-12, maIFNα or Alfacon (20 µL, in 2% FCS medium) was added to the cells directly after seeding. Supernatant (20 µL) was collected after 24 h and added to 180 µL Quanti-Blue^TM^ Solution (Invivogen) and incubated for 1 h (HEK-Blue™ ISG cells) or 4 h (B16-Blue™ IFN-α/β cells) at 37 °C after which the absorbance was measured at 630 nm.

Expression of mNG in BHK-ISRE-mNG cells following hsIFNα-2a, mmIFNα-1, maIFNα or Alfacon was determined as described above. Expression of mNG and miRFP670 was determined using a FACS Canto^TM^ II (BD Biosciences). Data were processed in FlowJo (for gating strategy see **Supplementary Figure S2C**). The EC_50_ values of each IFNα on each reporter cell line were calculated based on the predicted maximal values of that system, using a nonlinear regression model (least squares regression).

### Animals

All procedures involving experimental animals were approved by the relevant institutional animal ethics committees, conducted in accordance with applicable national regulations and FELASA guidelines, and reported herein in accordance with ARRIVE (Animal Research: Reporting of In Vivo Experiments) guidelines. Sample sizes were defined a priori by performing a power analysis with the expected effect size, statistical power of 80% and a significance level of 0.05, using G*Power3.1. Prior to experimental procedures, assessments focused on general health and welfare, including body condition, posture, grooming behavior and activity. During experimental procedures, for the development of clinical signs or adverse events.

Wild-type (WT) Syrian hamsters (*Mesocricetus auratus*; strain RjHan:AURA, male, 10-13 weeks old) were obtained specific pathogen-free from Janvier Laboratories (https://janvier-labs.com/en/, Le Genest Saint Isle, France) and allowed to acclimatise for one week prior to starting the experiment. Age- and sex-matched STAT2-KO hamsters were generated and bred in-house as previously described (19). Animals were housed and bred in individually-ventilated cages (maximum of 3 hamsters per cage, IsoCage N Biocontainment System, Tecniplast, Buguggiate, Italy) with a 12-h light–dark cycle (room temperature: 22 °C ± 2, humidity: 45–70%). Food and water were provided *ad libitum*, with cage enrichment (wood blocks) to encourage normal behavior. Bedding material was renewed weekly. Assignment of the groups was not blinded for operators. All hamster studies were conducted at the Rega Institute, KU Leuven. Experimental procedures and housing conditions were approved by the KU Leuven Ethical Committee (license DMIT-P116-2025) and conducted following Federation of European Laboratory Animal Science Associations (FELASA) guidelines.

### *In vivo* characterization of maIFNα

WT and STAT2-KO Syrian hamsters were randomly allocated to treatment or control groups. Animals in treatment groups received a single intraperitoneal (i.p.) dose of 5 µg/kg of purified maIFNα (100 µL in MEM medium containing 2% FCS). WT hamsters were euthanized (by i.p. injection of Dolethal, 500 µL; 200 mg/mL sodium pentobarbital, Vétoquinol SA, Magny-Vernois, France) at 2, 4, 6, 8, 18 or 24 h post-treatment and STAT2-KO animals at 6 h post-treatment after which the lung, liver, spleen and blood were collected. Non-treated animals (WT and STAT2-KO) were used as control groups. Organs were homogenized using bead disruption (innuSPEED Lysis Tube E, iST Innuscreen GmbH, Berlin, Germany) in TRK lysis buffer (E.Z.N.A.® Total RNA Kit, Omega Bio-tek, Pinnacle Way, USA) and centrifuged (16,000 × *g*, room temperature, 5 min) to pellet the cell debris. Blood was collected in TRK lysis buffer in Eppendorf tubes and centrifuged (16,000 × *g*, room temperature, 5 min). RNA was extracted following the manufacturer’s protocol and eluted in 50 µL. RNA levels of *MX2, ISG15* and *β-actin* were determined using the iTaq Universal Probes One-Step Kit (Biorad) on the LightCycler 96 Instrument (Roche) as described above.

### Statistics

All statistical analyses and data representation were performed using GraphPad Prism (version 10.6.1; GraphPad Software, Inc.). Dunnett’s multiple comparisons test (simple effects within columns or comparing cell means across rows and columns) was used to determine significance of ISG expression. EC_50_ and IC_50_ values were calculated by nonlinear regression (least squares fit) with fixed maximal values. Unpaired *t*-test with Welch’s correction was used to determine significant differences between 2 groups. Statistical significance was defined at *p* < 0.05.

## RESULTS

### Predicted maIFNα proteins are similar in primary sequence and structure to other IFNs

In contrast to many other species and despite its growing importance as preclinical animal model, the type I IFN gene cluster of the Syrian hamster is annotated solely based on prediction (25), void of functional confirmation. To assess whether these sequences encode functional IFNα proteins, we compared the predicted maIFNα (maIFNα-12-like) and maIFNα-9-like protein sequences with IFNα proteins from a range of vertebrate species.

Phylogenetic analysis placed the predicted maIFNα and maIFNα-9-like proteins within the rodent IFNα clade, together with *Mus musculus* (mm, house mouse) and *Rattus norvegicus* (rr, brown rat) sequences, with the closest relationship to IFNα proteins of other hamster species (*Cricetulus griseus*, cg, Chinese hamster; *Phodopus roborovskii*, pr, desert hamster). Both were more distantly related to *Homo sapiens* (hs, human) IFNα-2a, Alfacon, *Gallus gallus* (gg, chicken) IFNα, and *Danio rerio* (dr, zebrafish) IFN (**Figure 1A**).

**Figure 1.**
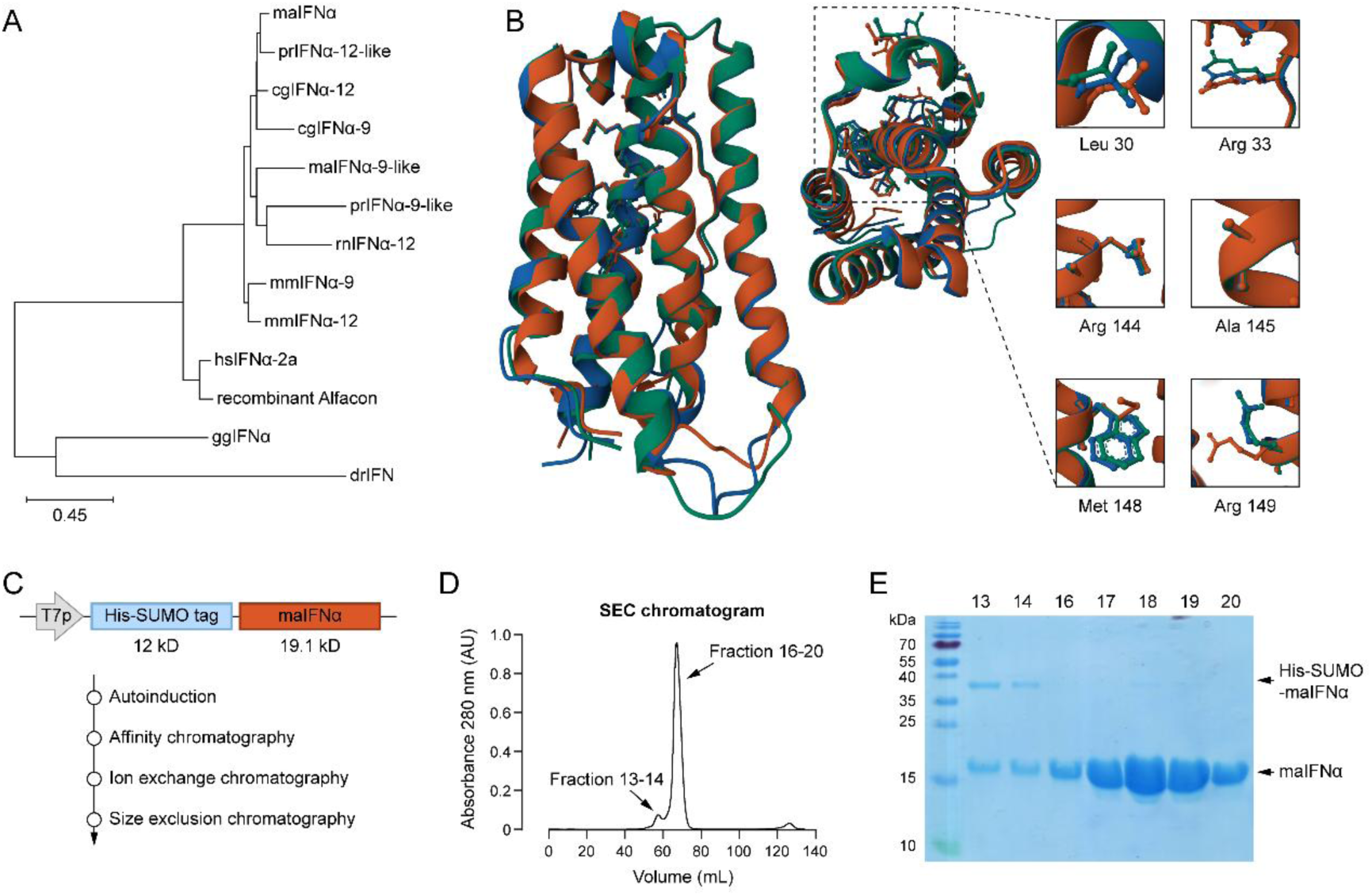
Comparison of maIFNα with other IFNs and subsequent protein purification. (A) Phylogentic tree of different IFNα proteins (without signal peptide) from different species, constructed via the Neighbor-Joining method. Scale bar represents AA substitutions per site. (B) Alphafold predicted protein structure of maIFNα (green), compared to hsIFNα-2a (red) and mmIFNα-12 (blue). Amino acids essential for binding of hsIFNα-2a to human IFNAR2 are indicated on the side view (left) and top view (middle). Inserts (right) show the position of the essential AA (in humans (3)), with corresponding AA from maIFNα and mmIFNα-12. (**C**) Schematic representation of the construct used for bacterial expression of maIFNα and overview of the different purification steps. (**D**) Chromatogram depicting the UV absorbance (280 nm) during SEC. Collected fractions are indicated with arrows. (**E**) SDS-PAGE of collected fractions following SEC. Fractions 16-20 were pooled and used for all subsequent experiments.

Amino acid sequence identity for maIFNα-9-like and maIFNα against cgIFNα-9, cgIFNα-12, prIFNα-12, mmIFNα-9 and mmIFNα-12 exceeded 63% and 78%, respectively, whereas identity to hsIFNα-2a reached at most 48% and 60%. Multiple sequence alignment showed comparable sequence lengths for maIFNα-9-like and maIFNα (159 and 167 AA, respectively) and the other IFNα proteins examined (160-167 AA) with several residues conserved across all species (**Supplementary Figure S1A**).

The structure of maIFNα was modelled using AlphaFold and aligned to hsIFNα-2a (pdb_00001rh2) and mmIFNα-12 (Q80SS5). TM-scores (43) of 0.94 and 0.95, respectively, indicate close structural correspondence between the predicted maIFNα model and both reference structures (**Figure 1B**). Together, these analyses indicate that the Syrian hamster type I IFNα gene cluster contains coding sequences with features expected for functional IFNα proteins.

### Expression and purification of predicted maIFNα protein

To obtain recombinant protein for functional characterization, the predicted maIFNα coding sequence was cloned into the pETSUK2 expression vector, downstream of a T7 promotor and a His_6_-SUMO tag (**Figure 1C**). The protein was expressed in *E. coli* Rosetta 2 (DE3) pLysS and purified by successive affinity, ion-exchange and size-exclusion chromatography (SEC), to a final yield of 21.5 mg/L (**Figure 1C**). The dominant main peak of the final SEC run (**Figure 1D**) reveals a >95% pure protein as judged by SDS-PAGE (**Figure 1E**), with the apparent mass closely matching the predicted molecular weight of maIFNα (19.1 kDa).

### MaIFNα induces ISGs through the JAK/STAT signaling pathway *in vitro*

To determine whether maIFNα retains its biological activity following expression in *E. coli* and purification, we evaluated its capacity to induce ISGs *in vitro*. BHK21J cells were stimulated with increasing concentrations (0-1,000 pg/mL) of maIFNα and the expression of *MX2* and *ISG15,* both canonical downstream targets of type I IFN signaling, was quantified by RT-qPCR. Gene expression levels were normalized to *β-ACTIN* and compared to unstimulated controls. While not detectable in unstimulated cells (Ct values > 40), *MX2* expression became detectable already after stimulation with as little as 1 pg/mL maIFNα (1.16 ± 0.19 log_10_-fold; *p* = 0.4444) and showed a dose-dependent increase reaching a 5.13 ± 0.10 log_10_-fold induction with 1,000 pg/mL maIFNα (*p* = 0.0108), with a resulting EC_50_ of 7.91 pg/mL (95% CI [5.43-11.49]). *ISG15* showed some basic expression (Ct values < 25) and was not further induced by 1-10 pg/mL maIFNα (-0.04 ± 0.04 log_10_-fold; *p* = 0.2989 and 0.13 ± 0.08 log_10_-fold; *p* = 0.5731) yet increased to 2.05 ± 0.07 log_10_-fold with 1,000 pg/mL maIFNα (*p* = 0.0015) (**Figure 2A**), with a resulting EC_50_ of 56.65 pg/mL (95% CI [43.01-74.96]). BHK21J cells were stimulated in parallel with Alfacon as a benchmark. Alfacon also induced both genes dose-dependently, *i.e.*, 1.93 ± 0.04 log_10_-fold for *MX2* (EC_50_ = 188.2 pg/mL; 95% CI [66.52-593.9]) and 2.16 ± 0.06 log_10_-fold for *ISG15* (EC_50_ = 382.5 pg/mL; 95% CI [233.6-474.5]) after induction with 1,000 pg/mL Alfacon (both *p* ≤ 0.0001) (**Figure 2A**). Hence particularly regarding *MX2* induction, maIFNα was markedly more potent than Alfacon on hamster cells, considering both the concentration needed for induction (about 24 times lower EC_50_; *p* ≤ 0.0001, Extra sum-of-squares F test) and the overall amplitude that could be reached (approximately 1500-times higher; *p* = 0.019, unpaired *t*-test with Welch’s correction).

**Figure 2.**
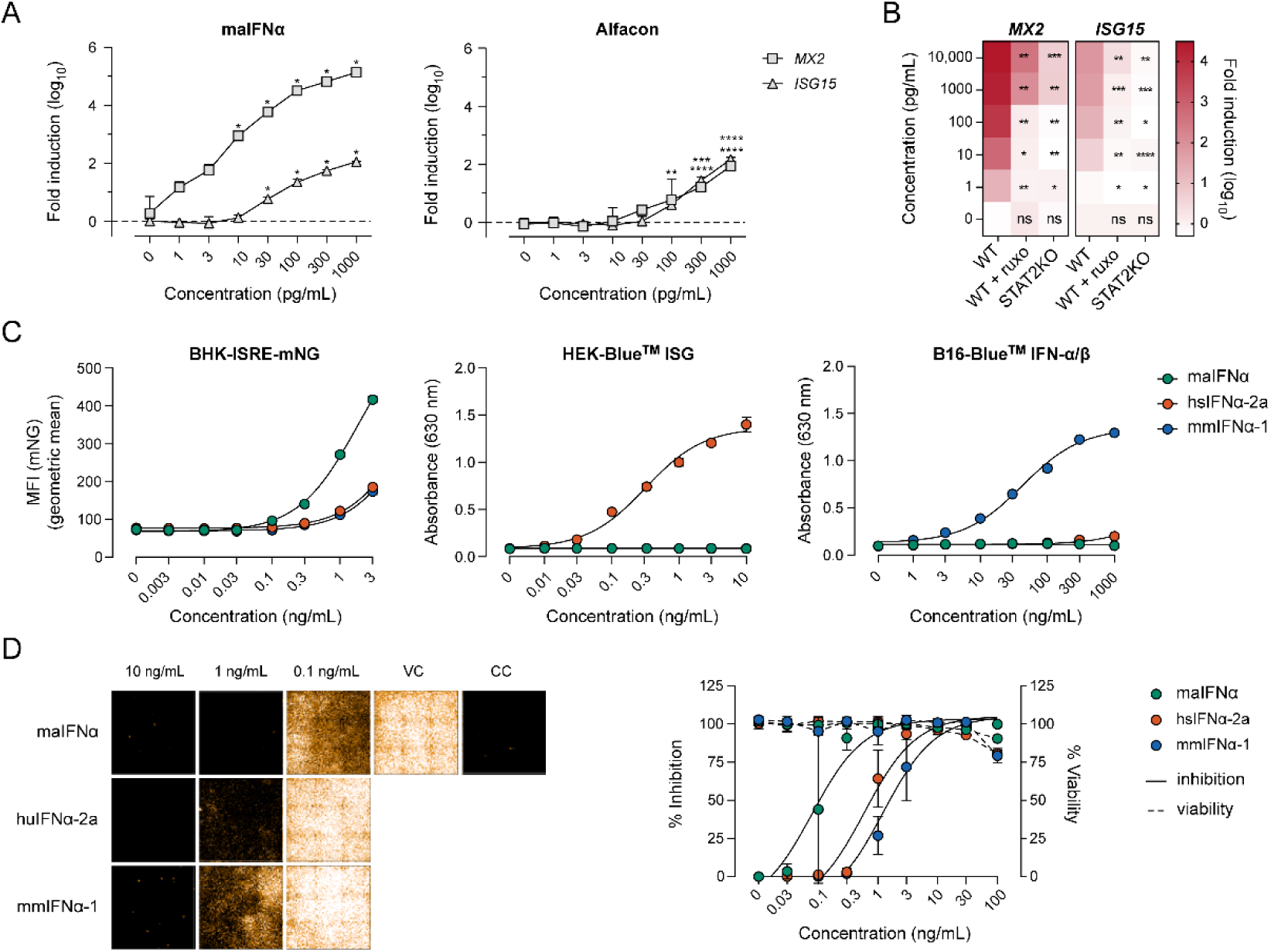
*In vitro* characterization of maIFNα. (**A**) Expression (log_10_-fold induction) of *MX2* and *ISG15* in BHK21J cells following maIFNα (left) or Alfacon (right) stimulation. Data points represent the mean ± SD of technical triplicates. Dunnett’s multiple comparison test, within group comparison. In the Alfacon graph: upper asterixis belong to *MX2*, lower asterixis belong to *ISG15*. (**B**) Heatmap showing expression (log_10_-fold induction) of *MX2* and *ISG15* in primary hamster fibroblasts cells (WT, WT treated with 5 µM ruxolitinib or STAT2-KO cells) following maIFNα stimulation (compared to unstimulated cells). Data points represent the median of technical triplicates. Dunnett’s multiple comparison test, within and between group comparison. (**C**) Species specificity of maIFNα, hsIFNα-2a and mmIFNα-2 on BHK-ISRE-mNG (mNG reporter), HEK-Blue^TM^ ISG (SEAP reporter) and B16-Blue^TM^ IFN-α/β (SEAP reporter) cells. Data points represent the mean ± SD of technical triplicates. EC_50_ values were calculated by nonlinear regression (least squares fit) with fixed maximal values. (**D**) Antiviral activity of maIFNα, hsIFNα-2a and mmIFNα-2 on BHK-miRFP670 cells inoculated with YF17D-mCherry reporter virus, at 6 dpi. Representative pictures (complete well) of infection (left) and % inhibition of viral replication and % cell viability (right). Data points represent the mean ± SD of technical triplicates. IC_50_ values were calculated by nonlinear regression (least squares fit) with fixed maximal values. P-values: ns – p > 0.05; * – p ≤ 0.05; ** – p ≤ 0.01; *** – p ≤ 0.001; **** – p ≤ 0.0001.

MaIFNα activity was subsequently established in primary WT Syrian hamster fibroblasts. Following stimulation with 1-10,000 pg/mL maIFNα, *MX2* expression increased from 1.29 ± 0.17 (*p* = 0.0136) to 4.49 ± 0.15 log_10_-fold (*p* = 0.0016), whereas *ISG15* expression increased from -0.11 ± 0.18 (*p* = 0.9795) to 1.87 ± 0.09 log_10_-fold (*p* = 0.0080) (**Figure 2B**) with EC_50_ values of 3.87 ng/mL (95% CI [1.29-8.37]) for *MX2* and of 19.24 ng/mL (95% CI [11.60-33.89]) for *ISG15*, respectively. Alfacon induced comparable levels of both transcripts (**Supplementary Figure S2B**).

To confirm whether that ISG induction was mediated by the canonical JAK/STAT signaling pathway, primary WT Syrian hamster fibroblasts were pre-treated with the JAK1/JAK2 inhibitor ruxolitinib (5 µM) 1 h before stimulation with maIFNα. Under these conditions, *MX2* induction was largely suppressed and detected only following stimulation with 1,000 and 10,000 pg/mL maIFNα (2.03 ± 0.16 log_10_-fold; *p* = 0.0225 and 2.56 ± 0.05 log_10_-fold; *p* = 0.0260), and transcript levels were significantly lower than in non-treated WT fibroblasts stimulated with 10,000 pg/mL maIFNα (*p* = 0.0016) (**Figure 2B**). Also *ISG15* expression was largely inhibited and showed a significant induction only after stimulation with a as high concertation as 10,000 pg/mL maIFNα relative to unstimulated (but treated) controls (*p* = 0.0008), again at significantly lower levels of transcripts than in non-treated WT fibroblasts stimulated at the same concentration (*p* = 0.0025) (**Figure 2B**).

The same approach was applied to primary STAT2-KO Syrian hamster fibroblasts. Indeed STAT2-KO fibroblasts appeared more than 10^6^ times more refractory to maIFNα stimulation than WT fibroblasts, whereby some limited *MX2* expression became detectable only after stimulation at the highest applied concentrations (1,000 pg/mL: 0.93 ± 0.43 log_10_-fold; *p* = 0.0060 and 10,000 pg/mL: 0.69 ± 0.04 log_10_-fold; *p* = 0.2633), and failed to induce any measurable *ISG15* expression at any concentration tested (**Figure 2B**). Likewise, when compared to primary WT fibroblasts, the transcript levels of both genes were significantly lower (*p* = 0.0006 and *p* = 0.0072, respectively). Comparable effects were observed in control experiments using Alfacon stimulation of ruxolitinib-treated primary WT and STAT2-KO Syrian hamster fibroblasts (**Supplementary Figure S2B**). Together, these results indicate that the predicted maIFNα protein is biologically active and induces ISG expression through the canonical JAK/STAT signaling pathway.

### MaIFNα displays species-restricted activity

The species specificity of maIFNα, hsIFNα-2a and mmIFNα-1 was determined by comparing their relative activities using reporter cell lines derived from hamster (BHK-ISRE-mNG; mNG reporter), human (HEK-Blue™ ISG; SEAP reporter) and mouse (B16-Blue™ IFN-α/β; SEAP reporter). In BHK-ISRE-mNG cells we used flow to quantify the mean fluorescent intensity (MFI) of mNG following stimulation with IFNs. In the HEK-Blue™ ISG and B16-Blue™ IFN-α/β cells, we measured the absorbance (630 nm) to determine the activity of the SEAP reporter protein in the supernatant of stimulated cells.

The BHK-ISRE-mNG reporter was first validated by stimulation with a serial dilution of maIFNα or Alfacon, which induced significant mNG expression relative to unstimulated cells. As expected, expression of the nuclear miRFP670 protein, which is under a constitutive EFIα promotor, remained stable (**Supplementary Figure S2D**).

Next, each IFN was tested across serial dilutions, using concentration ranges optimized for each reporter system. MaIFNα was a potent inducer of mNG in BHK-ISRE-mNG cells (EC_50_ = 1.81 ng/mL; 95% CI [1.73-1.88]), but showed no detectable activity in HEK-Blue™ ISG or B16-Blue™ IFN-α/β cells at the concentrations tested. HsIFNα-2a and mmIFNα-1 were also able to induce reporter protein expression in BHK-ISRE-mNG cells, though with lower potency (EC_50_ = 12.03; 95% CI [11.54-12.54] and EC_50_ = 12.94; 95% CI [12.28-13.66], respectively) (Figure 2C, Table 3).

**Table 3.** EC_50_ values and confidence intervals of different IFNs on reporter cell lines.

|  | **BHK-ISRE-mNG<br>EC<sub>50</sub><br>[95% CI]** | **HEK-Blue™ ISG<br>EC<sub>50</sub><br>[95% CI]** | **B16-Blue™ IFN-α/β<br>EC<sub>50</sub><br>[95% CI]** |
|---|---|---|---|
| maIFNα | 1.805 ng/mL<br>[1.734-1.881] | na | na |
| hsIFNα-2a | 12.03 ng/mL<br>[11.54-12.54] | 0.3164 ng/mL<br>[0.2635-0.3805] | na |
| mmIFNα-1 | 12.94 ng/mL<br>[12.28-13.66] | na | 42.58 ng/mL<br>[37.79-48.02] |
| Alfacon | 90.61 ng/mL<br>[83.96-97.84] | 8.025 ng/mL<br>[6.332-10.19] | 61.86 ng/mL<br>[51.51-74.48] |

na – not active

HsIFNα-2a was most active on HEK-Blue™ ISG cells (EC_50_ = 0.3164 ng/mL; 95% CI [0.2635-0.3805]), and inactive on B16-Blue™ IFN-α/β cells, whereas mmIFNα-1 was most active on B16-Blue™ IFN-α/β cells (EC_50_ = 42.58 ng/mL; 95% CI [37.79-48.02]) and inactive on HEK-Blue™ ISG cells (**Figure 2C**, **Table 3**). Alfacon, which is considered to be a universally active IFN, was included as a positive control at identical concentrations in all three reporter cell lines (**Supplementary Figure S2E,** **Table 3**). In summary, each IFN exhibited its highest biological activity on the homologous system, while also demonstrating a limited heterologous cross-species activity.

### Pronounced antiviral effect of maIFNα

The antiviral activity of maIFNα, hsIFNα-2a and mmIFNα-1 was evaluated in BHK-miRFP670 cells inoculated with a YF17D-mCherry reporter virus. Cells were pre-treated with each IFN for 1 h prior to inoculation (MOI 0.01), and viral replication was assessed at 6 dpi based on mCherry reporter expression. All three IFNs inhibited the viral replication in a dose-dependent manner, as demonstrated by loss of the mCherry signal with increasing IFN concentrations (**Figure 2D**). MaIFNα showed the greatest antiviral potency on hamster cells (IC_50_ = 0.079 ng/mL; 95% CI [0.051-0.12]), followed by hsIFNα-2a (IC_50_ = 0.59 ng/mL; 95% CI [0.41-0.85]) and mmIFNα-12 (IC_50_ = 1.23 ng/mL; 95% CI [0.80-1.83]). Alfacon, included as a positive control, produced a comparably dose-dependent yet less pronounced inhibitory effect (IC_50_ = 6.016 ng/mL; 95% CI [3.95-9.25]) (**Supplementary Figure S2F**). The cytostatic activity of IFNα is expected to interfere with cell viability. Cytotoxicity was hence assessed in parallel on BHK-miRFP670 cells. Limited toxicity was observed only at the highest concentration tested (100 ng/mL), with cell viability of 80.6% (*p* = 0.0336) for maIFNα, 79.3% (*p* = 0.0157) for hsIFNα-2a and 90.6% (*p* = 0.551) for mmIFNα-1. Therefore, as demonstrated for YF17D, homologous maIFNα inhibits viral replication in hamster cells with greater potency than either heterologous hsIFNα-2a or mmIFNα-1 (**Figure 2D**).

### Purified maIFNα induces ISG expression *in vivo*

The capacity of purified maIFNα to induce ISG expression *in vivo* was evaluated in WT Syrian hamsters. To concomitantly monitor its pharmacodynamics, animals (10-13 weeks old) received a single administration of maIFNα (5 µg/kg, i.p.) and were euthanized at 2, 4, 6, 8, 18 or 24 h post-treatment (hpt), after which *MX2* and *ISG15* expression were determined in the blood, spleen, liver and lung (**Figure 3A**).

**Figure 3.**
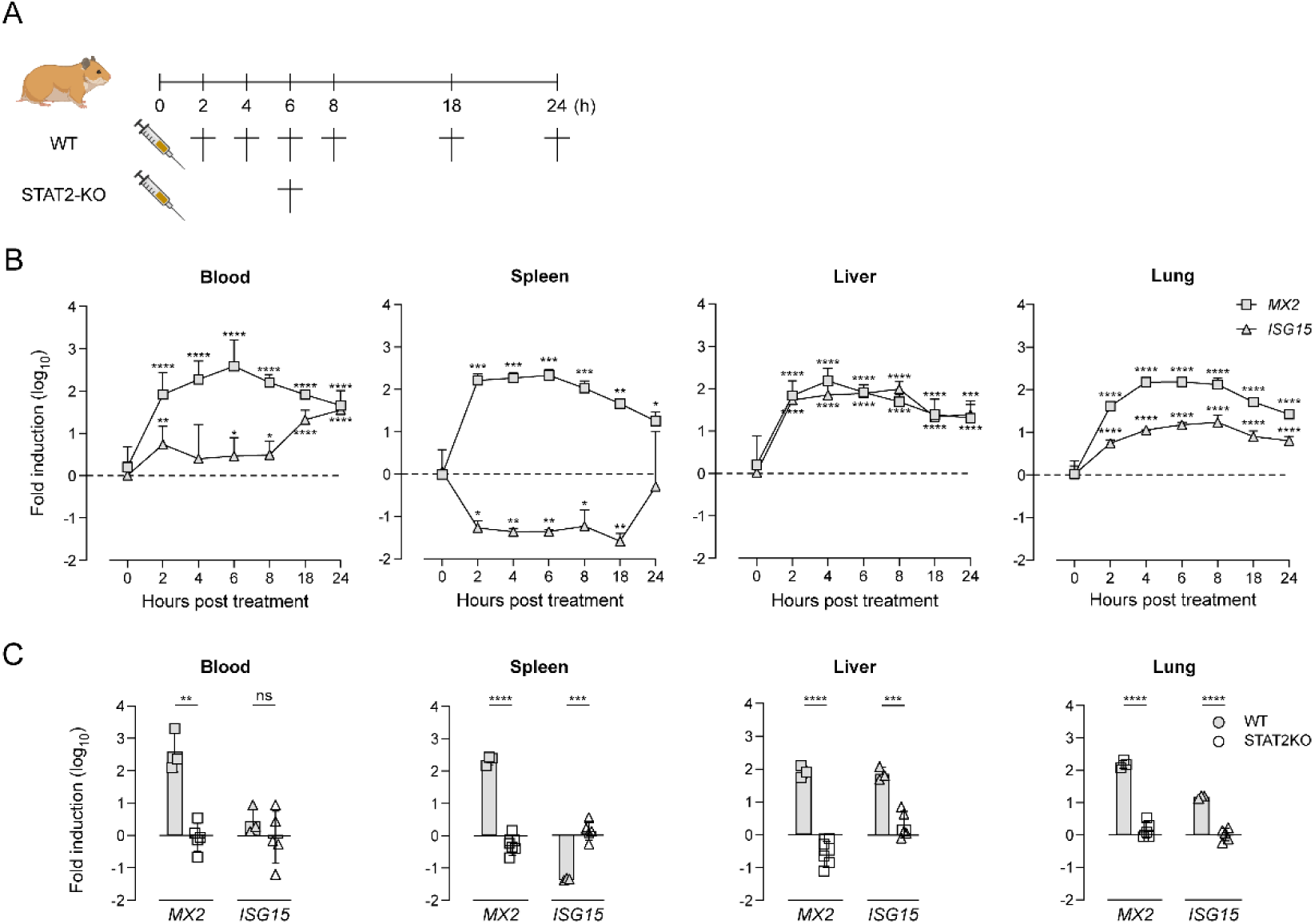
Kinetics of maIFNα *in vivo*. (**A**) Schematic representation of maIFNα treatment in WT and STAT2-KO hamsters. (**B**) Kinetics of *MX2* and *ISG15* expression in blood, spleen liver and lung in WT hamsters (n = 3 per group). Data points represent the mean ± SD. Dunnett’s multiple comparison test, within group comparison. (**C**) Expression (log_10_-fold) of *MX2* and *ISG15* in blood, spleen, liver and lung in STAT2-KO hamsters (n = 5) compared to WT hamsters (n = 3). Unpaired *t*-test with Welch’s correction. P-values: ns – p > 0.05; * – p ≤ 0.05; ** – p ≤ 0.01; *** – p ≤ 0.001; **** – p ≤ 0.0001.

*MX2* was induced in all tissues examined, with significant induction already detected at 2 hpt in blood (1.92 ± 0.51 log_10_-fold; *p* = 0.001), spleen (2.21 ± 0.15 log_10_-fold; *p* = <0.0001), liver (1.83 ± 0.35 log_10_-fold; *p* = 0.0005) and lung (1.91 ± 0.088 log_10_-fold; *p* = <0.0001) relative to untreated controls (0 hpt). Expression peaked at 6 hpt in blood (2.58 ± 0.62 log_10_-fold; *p* = <0.0001), spleen (2.33 ± 0.14 log_10_-fold; *p* = <0.0001) and lung (2.18 ± 0.12 log_10_-fold; *p* = <0.0001), and declined thereafter, whereas in the liver the maximum was reached at 4 hpt (2.18 ± 0.29 log_10_-fold; *p* = <0.0001) (**Figure 3B**). *ISG15* followed different kinetics. In blood, a significant induction was detected only from 18 hpt (1.32 ± 0.23 log_10_-fold; *p* = 0.0108), reaching a maximum at 24 hpt (1.55 ± 0.16 log_10_-fold; *p* = 0.0031). In the spleen, *ISG15* expression was downregulated at 2 hpt relative to untreated controls (-1.27 ± 0.17 log_10_-fold; *p* = 0.0473). In liver and lung, induction was already significant at 2 hpt (1.73 ± 0.085 log_10_-fold; *p* = <0.0001 and 0.74 ± 0.082 log_10_-fold; *p* = 0.0002) and peaked at 8 hpt (1.99 ± 0.18 log_10_-fold; *p* = <0.0001 and 1.23 ± 0.17; *p* = <0.0001) (**Figure 3B**).

To establish that this induction was mediated by the canonical type I IFN-induced JAK/STAT signaling rather than by confounding factors, STAT2-KO hamsters (10-13 weeks old) received maIFNα (5 µg/kg, i.p.) and were euthanized 6 hpt. Neither *MX2* nor *ISG15* was induced in blood (-0.05 ± 0.43 and -0.29 ± 1.31 log_10_-fold), liver (-0.65 ± 0.33 and 0.32 ± 0.41 log_10_-fold), spleen (-0.30 ± 0.30 and 0.14 ± 0.29 log_10_-fold) or lung (0.16 ± 0.25 and 0.01 ± 0.19 log_10_-fold). Expression differed significantly between WT and STAT2-KO animals for both genes in spleen (*p* = <0.0001 and *p* = 0.0003), liver (*p* = <0.0001 and *p* = 0.0004) and lung (*p* = <0.0001 and *p* = <0.0001), and for *MX2* in blood (*p* = 0.0064) (**Figure 3C**). These results indicate that purified maIFNα is able to induce a systemic ISG response *in vivo,* and that this response depends on canonical STAT2-mediated type I IFN signaling.

## Discussion

Although Syrian hamsters constitute an increasingly valued model to study infectious diseases, the structure and activity of hamster IFNα as a key player orchestrating antiviral responses remains poorly described. In this study, we report the production of predicted maIFNα by recombinant expression and its functional validation. We document the species-specific activity of maIFNα to induce canonical type I IFN responses, measured as ISG induction *in vitro* and *in vivo* and potent antiviral activity when tested on hamster cells, alike that observed for the IFNα proteins of other species (44,45).

The chromosomal location of the maIFNα gene cluster has not yet been identified, however more insights were obtained from hamster genome assembly MesAur 2.0 (25), indicating 14 predicted IFNα genes and 5 pseudogenes. This gene cluster is flanked by *IFNB* and *IFNE* genes, showing a very similar genomic organization to that mapped in mice (46). We selected one of the 14 predicted genes encoding for maIFNα-12-like (XP_040613374.1) as representative maIFNα based on bioinformatical analysis (**Figure 1**, **Supplementary Figure S1**) for in-depth functional characterization. The maIFNα sequence showed highest sequence similarity to the respective orthologs from Chinese (XP_027256148.1) and dwarf hamsters (XP_051043952.1). The Alphafold model of maIFN showed an almost perfect fit (TM-scores of 0.94 and 0.95) with the crystal structures of hsIFNα-2a (pdb_00001rh2) and mmIFNα-12 (Q80SS5) with key structural elements such as the overall four α-helices scaffold conserved, while differences in AA residues shown to be key for high-affinity IFNAR2 binding in the human system (47) (**Figure 1B**, **Supplementary Figure S1A**) may explain its superior activity on hamster cells.

Glycosylation of type 1 IFNs has been shown to increase IFN serum stability and half-life (48) and endogenous maIFNα is possibly glycosylated at position 135 (**Supplementary Figure S1A**) (23,24), while IFNs expressed in *E. coli* lack such post-translational modifications (31–33,49,50). Seen the high specific activity observed for maIFNα (EC_50_ values of about 0.25 -2.5 pM), glycosylation appears dispensable. In line, with exception of hsIFNα-2 (51) and hsIFNα-14 (52), most human type I IFNα subtypes are unglycosylated. Also, while the majority of mouse IFNα subtypes contains putative N-glycosylation sites no correlation was found with activity (53). Nevertheless, here we show that glycosylation is not required for the biological function (*in vitro* and *in vivo*) of recombinant maIFNα, despite the presence of a putative N-glycosylation site.

Species-specificity appears to be more relevant than potential post-translational modifications. Previous studies report varying *in vitro* cross-species activities of different human IFNα subtypes on selected cell types of other species (54–56), whereby recombinant human IFNα

Alfacon (Universal^TM^ Type I IFN) seems to have a fairly broad host-range *in vitro* (57–59) and *in vivo*, demonstrated for instance in calves (60), cats (61), mice (58) and hamsters (38,62,63). Data reporting the *in vitro* activity across species for mouse IFNα subtypes remain conflicting (64–68). We demonstrate a rather restricted cross-species activity of maIFNα, hsIFNα-2a and mmIFNα-1, at least in the range of herein tested *in vitro* systems (hamster cell lines and primary hamster fibroblasts; reporter cells originating from different host species), whereby maIFNα showed a markedly higher potency than its human and murine orthologs in the hamster system (**Figure 2**, **Table 3**).

The absolute potency of different IFNs is difficult to compare, particularly for orthologs from other species, as no universally accepted reference standard analogous to the WHO International Standard for human IFNα, which defines International Units (IU), is available (69). Alfacon, originally commercialized as Infergen® (Astellas Pharma; EMEA/H/C/000186) for the treatment of chronic hepatitis C, was standardized to a biological potency of approximately 10⁹ IU/mg protein. However, its measured antiviral activity varies widely depending on the virus species and particular cell-based assay used. Notably, the antiviral activity of maIFNα against YF17D in BHK21 cells, with an IC_50_ of approximately 0.1 ng/mL (about 5 pM) (**Figure 2D**), closely matches that reported for Alfacon against YF17D in African green monkey (Vero) cells (70).

*In vitro*, IFNα is known to trigger a rapid induction of ISGs followed by their downregulation through negative-feedback mechanisms. Despite the widespread clinical use of IFNα for the treatment of viral infections (71) and malignancies (72), surprisingly little is known about the pharmacodynamics of single-dose administration *in vivo*. However, available data from murine models suggest that the transient response observed *in vitro* is similarly reproduced *in vivo* (73,74). Accordingly, systemic administration of maIFNα leads to a robust yet transient upregulation of ISGs, represented by rapid changes in *MX2* and *ISG15* expression levels in the blood, spleen, liver and lung of hamsters in a STAT2-dependent manner, peaking within 4 to 6 hours after a single 5 µg/kg dose. Nevertheless, this fast immediate response started contracting within 24 h after dosing. The only exception was an inverse effect observed for *ISG15* in the spleen. This selective regulation of *ISG15* expression is consistent with an auto-regulatory negative-feedback loop, reflecting the increasingly recognized role of *ISG15* and the deISGylase *USP18* in limiting the duration and magnitude of IFN signaling (75,76).

Alfacon has previously been evaluated as antiviral treatment in Syrian hamsters *in vivo*. In addition to studies involving low-pathogenicity prototype viruses representing *Picornaviridae* (38), *Arenaviridae* (77), or *Phenuiviridae* (78), it has also been investigated for the experimental treatment of severe arboviral diseases in hamsters, including for yellow fever (79), West Nile fever (80,81) and Western equine encephalitis (63). However, therapeutic efficacy has remained limited, even at relatively high doses of up to 5 μg/kg/day in these animals. This contrasts with the substantially lower doses typically used in humans, where prolonged treatment with Alfacon corresponds to approximately 0.1-0.25 μg/kg/day (assuming a 15 μg daily dose). In lack of characterization of the thus-induced IFN response, reasons for failure remain elusive. Nevertheless, with the availability of maIFNα, it is now possible to directly assess the primary determinants of antiviral efficacy: whether the limited therapeutic benefit observed previously reflects the ability of the pathogens to evade innate immune responses through sophisticated IFN-antagonistic mechanisms (11,12), thereby restricting the efficacy of exogenous IFN therapy, or whether it is largely attributable to species-specific differences in IFNAR engagement and activation. The latter could be overcome by administering the homologous IFNα in the corresponding animal host.

In current study, we explored only one maIFNα subtype out of 14 maIFNα-12-like and 2 maIFNα-9-like alternative gene products. Further analysis of hamster IFNβ (maIFNβ; XM_040757177.1) was skipped as preliminary assessment of maIFNβ produced by transient transfection of HEK-AD cells suggest a lower potency compared to maIFNα (not shown). By analogy with humans (82) and mice (53), different IFNα subtypes can be expected to vary widely in their affinity for the common IFNAR, resulting in marked differences in their *in vitro* potency regarding kinetics and magnitude of ISG induction and associated biological activity. How this translates into different IFNs differentially orchestrating innate and adaptive immune responses *in vivo* remains poorly understood even in the best studied murine system. Overall, the discovery of maIFNα and the establishment of a validated toolbox to study innate IFN signaling in hamsters are further steps in unlocking experimental limitations in the use of Syrian hamsters as an increasingly valued infectious diseases model.

## ACKNOWLEDGMENTS

The authors are grateful to Thibault Francken for kindly providing BHK-miRFP670 cells and to Viktor Lemmens for providing the pLenti-EFIα-H2B-mi670-P2A-BlaR plasmid. The psPAX2 and pMD2.G were a gift from Didier Trono. The authors thank Geert Schoofs for help with the flow cytometry data acquisition and to Carolien De Keyzer, Lindsey Bervoets and the staff of the animal facility at the Rega Institute (KU Leuven) for excellent support during animal experiment, and Viktor Lemmens, Thibault Francken, Joost Schepers and Winston Chiu for their support with HCI imaging.

## CRediT Authorship Contribution Statement

Conceptualization, V.N., Y.A.A., K.D.

Data curation, V.N., S.B., Y.A.A.

Formal analysis, V.N., S.B., Y.A.A.

Funding acquisition, K.D.

Investigation, V.N., S.B., Y.A.A.

Methodology, V.N., S.B., S.S., E.B., Y.A.A., K.D.

Project administration, Y.A.A., K.D.

Resources, S.S, K.D.

Supervision, Y.A.A., K.D.

Validation, Y.A.A., K.D.

Visualization, V.N., Y.A.A.

Writing – original draft, V.N., Y.A.A., B.L., K.D.

Writing – review & editing, all authors.

All authors have read and agreed to the published version of the manuscript.

## Data and materials availability

All data supporting the findings in this study are available from the corresponding author upon reasonable request.

