## Supplementary Figures for "*In vitro* and *in vivo* validation of recombinant Syrian hamster interferon-α"

**Supplementary Figure 1**

**Supplementary Figure 2**

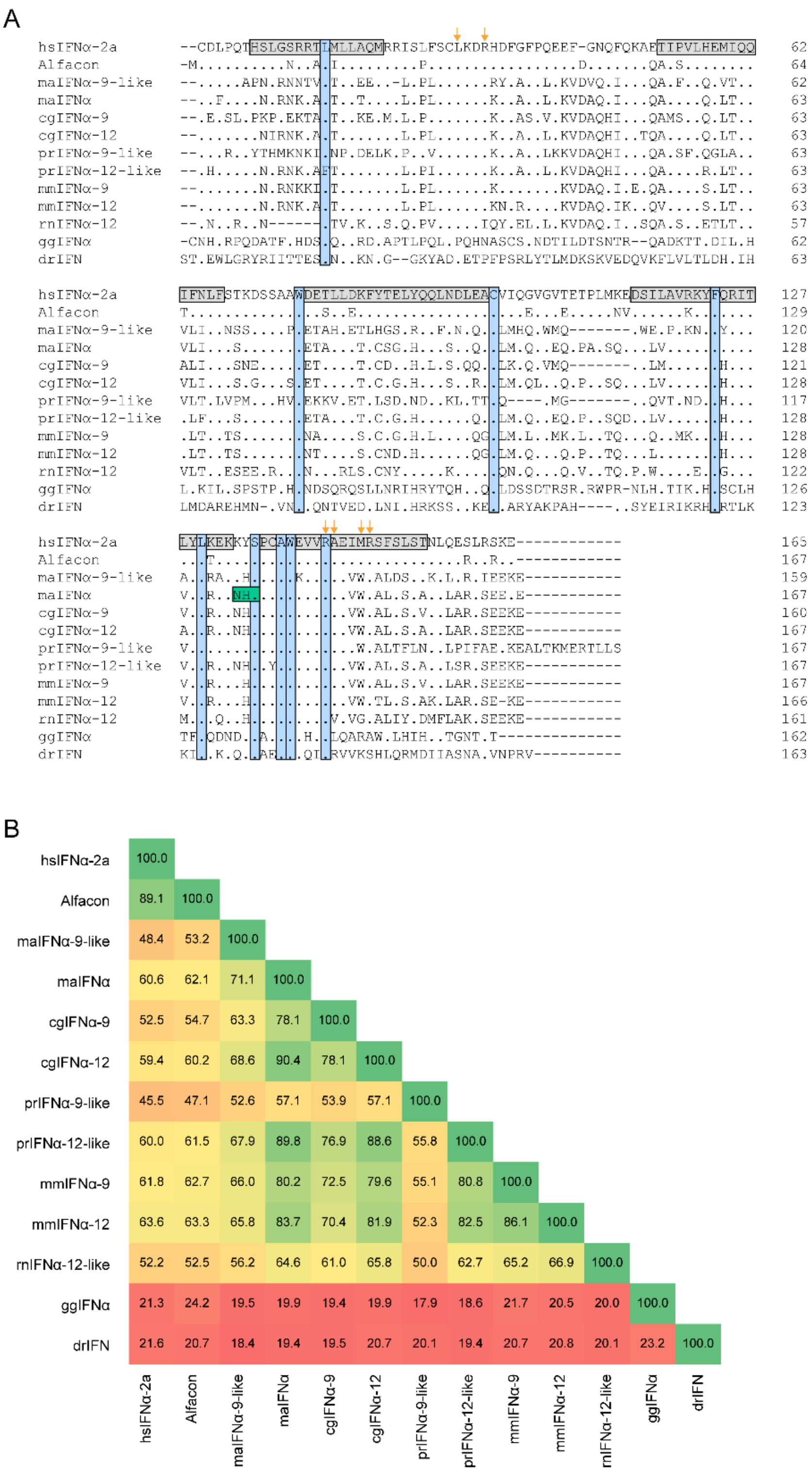


**Supplementary Figure 1. Sequence alignment and percent identity matrix.**

(**A**) Sequence alignment of IFNα proteins from different species. Signal peptide is excluded. Dots represent identical AA to huIFNα-2a. Dashes represent gaps in the sequence. Grey bars indicate the presence of a α-helix in the secondary protein structure of hsIFNα-2a. Blue bars indicate conserved AA across all sequences. Green bar indicates a putative N-glycosylation site of maIFNα. The orange arrows indicate essential AA in hsIFNα-2a for binding with human IFNAR2. (**B**) Percent identity matrix of IFNα proteins from different species. Signal peptide is excluded.

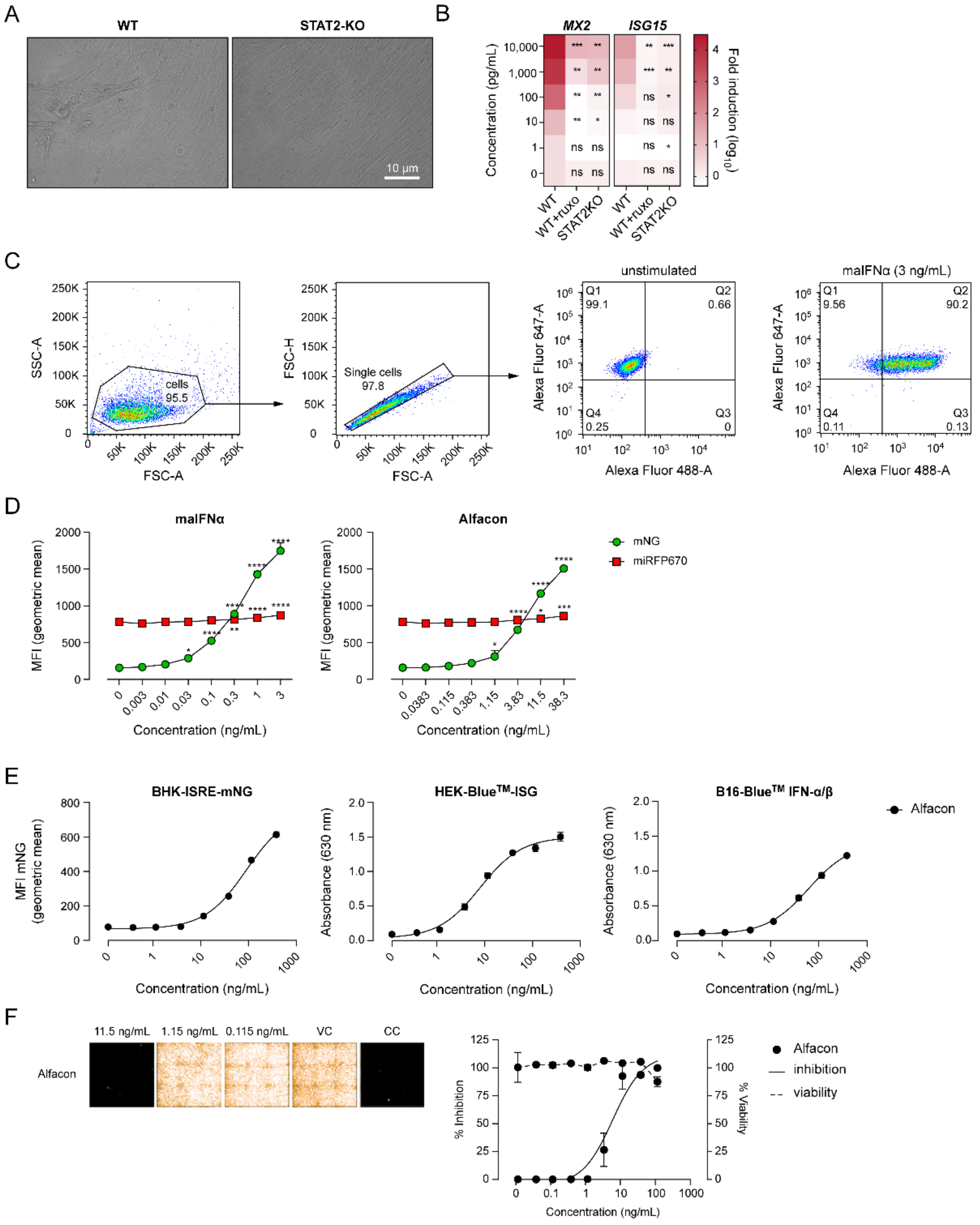


**Supplementary Figure 2. Alfacon as positive control for maIFNα**.

(**A**) Representative pictures of primary fibroblasts isolated from WT and STAT-2KO hamsters, 14 days post isolation. (**B**) Heatmap showing expression ($log_{10}$-fold induction) of *MX2* and *ISG15* in primary hamster fibroblasts cells (WT, WT treated with 5 µM ruxolitinib or STAT2-KO cells) following Alfacon stimulation (comparted to unstimulated cells). (**C**) Gating strategy applied following maIFNα stimulation on BHK-ISRE-mNG cells. (**D**) Validation of BHK-ISRE-mNG reporter cell line, showing expression of mNG and miRFP670 following stimulation with maIFNα (left) and Alfacon (right). (**E**) Alfacon stimulation on reporter cells derived from hamster (BHK-ISRE-mNG, mNG reporter), human (HEK-Blue™-ISG, SEAP reporter) and mouse (B16-Blue™ IFN-α/β, SEAP reporter) cells. Data points represent the mean ± SD of technical triplicates. (**F**) Antiviral activity of Alfacon on BHK-miRFP670 cells infected with YF17D-mCherry reporter virus at 6 dpi. Left panel, representative fluorescent pictures (9 pictures per well) of infected cells; right panel, inhibition and cell viability at indicated concentrations of Alfacon. Data points represent the mean ± SD of technical triplicates.
